# Fungal Induced Protein Hyperacetylation Identified by Acetylome Profiling

**DOI:** 10.1101/057174

**Authors:** Justin W Walley, Zhouxin Shen, Maxwell R. McReynolds, Steven P. Briggs

## Abstract

Lysine acetylation is a key post-translational modification that regulates diverse proteins involved in a range of biological processes. The role of histone acetylation in plant defense is well established and it is known that pathogen effector proteins encoding acetyltransferses can directly acetylate host proteins to alter immunity. However, it is unclear whether endogenous plant enzymes can modulate protein acetylation during an immune response. Here we investigate how the effector molecule HC-toxin, a histone deacetylase inhibitor, produced by *Cochliobolus carbonum* race 1 promotes pathogen virulence in maize through altering protein acetylation. Using mass spectrometry we globally quantified the abundance of 3,636 proteins and the levels of acetylation at 2,791 sites in maize plants treated with HC-toxin as well as HC-toxin deficient or producing strains of *C. carbonum.* Analyses of these data demonstrate that acetylation is a widespread post-translational modification impacting proteins encoded by many intensively studied maize genes. Furthermore, the application of exogenous HC-toxin enabled us to show that the activity of plant-encoded enzymes can be modulated to alter acetylation of non-histone proteins during an immune response. Collectively, these results provide a resource for further mechanistic studies examining the regulation of protein function and offer insight into the complex immune response triggered by virulent *C. carbonum.*

## INTRODUCTION

Protein lysine acetylation is an evolutionarily conserved reversible covalent modification. While lysine acetylation was discovered on histones more than 50 years ago it has long been known that non-histone proteins are also acetylated (1–4). Recently mass spectrometry based global acetylation profiling methods have been developed, leading to the realization that lysine acetylation is a major post-translational modification that impacts a wide-range of proteins (5–17). Acetylation of lysine residues is a covalent attachment typically added to or removed from proteins by histone acetyltransferase (HAT) and histone deacetylase (HDAC) enzymes, respectively (18). Alternatively, recent studies have demonstrated that protein acetylation is not only controlled enzymatically, but that it is also modulated non-enzymatically by metabolic intermediates including Acetyl-CoA and NAD^+^ (19, 20).

It is becoming increasingly apparent that modulation of protein acetylation plays a central role during host-pathogen interactions (21). Much of the research thus far has focused on how alterations in histone acetylation alter plant immunity. Modulation of histone acetylation was first implicated in plant immunity when it was discovered that the effector molecule HC-toxin, secreted by *Cochliobolus carbonum* race 1, functioned as a histone deacetylase inhibitor (HDACi) in fungi, plants, and animals (22–24).

Additionally, mutation or overexpression of HATs or HDACs in Arabidopsis and rice alters plant immunity and has been interpreted to be a direct result of modulating histone acetylation status and thus affecting transcription at specific defense gene promoters (25–27).

Pathogen infection also triggers alterations in host non-histone protein acetylation status and immunity (21, 28–32). To date pathogen induced non-histone protein acetylation has been shown to be a result of pathogen effector molecules functioning as acetyltransferase enzymes that act directly on host proteins. This raises the question of whether plant HATs or HDACs directly modulate acetylation status of non-histone proteins during pathogen infection.

In this study we used mass spectrometry to quantify global changes in protein abundance and acetylation levels triggered by pathogen infection. Critically, through the application of exogenous HC-toxin and *C. carbonum* strains we demonstrate that the activity of plant encoded enzymes can be modulated to alter both histone and non-histone protein acetylation. Additionally, we provide the first global acetylome for maize and significantly expand the number of acetylation sites identified in plants. Acetylated proteins carry out a wide-range of functions and include numerous well-studied maize proteins. Thus, the data presented here will enable new research approaches to understand the post-translational regulation and function of both defense and non-defense related proteins.

## RESSULTS

### Mapping the Maize Acetylome

We used mass spectrometry-based proteomics to globally identify acetylated proteins and uncover specific acetylation levels altered during pathogen infection. We profiled *Hm1A* mutant maize plants, which are susceptible to the fungal pathogen, *C. carbonum* race 1 (33). Plants were treated with 1) mock, 2) exogenous HC-toxin, 3) an HC-toxin deficient (Tox^-^) strain of *C. carbonum* and 4) HC-toxin producing (Tox^+^) strains of *C. carbonum* (Fig. 1A). To capture early signaling events we collected tissue at 22 hrs, which is before visual symptoms develop during a susceptible infection. These treatments were chosen to globally map protein acetylation sites in maize leaves and to measure changes induced by *C. carbonum* or HC-toxin.Total protein was extracted from the samples and tryptic peptides were labeled with iTRAQ reagents to quantify non-modified protein abundance (34). In parallel, we enriched acetylated peptides with an antibody that recognizes acetyl-lysine and then quantified the immunopurified peptides using spectral counting (5, 35). Using stringent cutoffs to maintain a low false discovery rate (0.1% peptide level FDR), we compared the levels of 3,636 non-modified proteins and 2,791 acetylation sites originating from 912 acetylated proteins (Fig. 1B and Datasets S1&S2). The 2,791 sites are more than double the number of previously reported acetylation sites and therefore significantly expand our knowledge of plant acetylomes.

**Figure 1.**
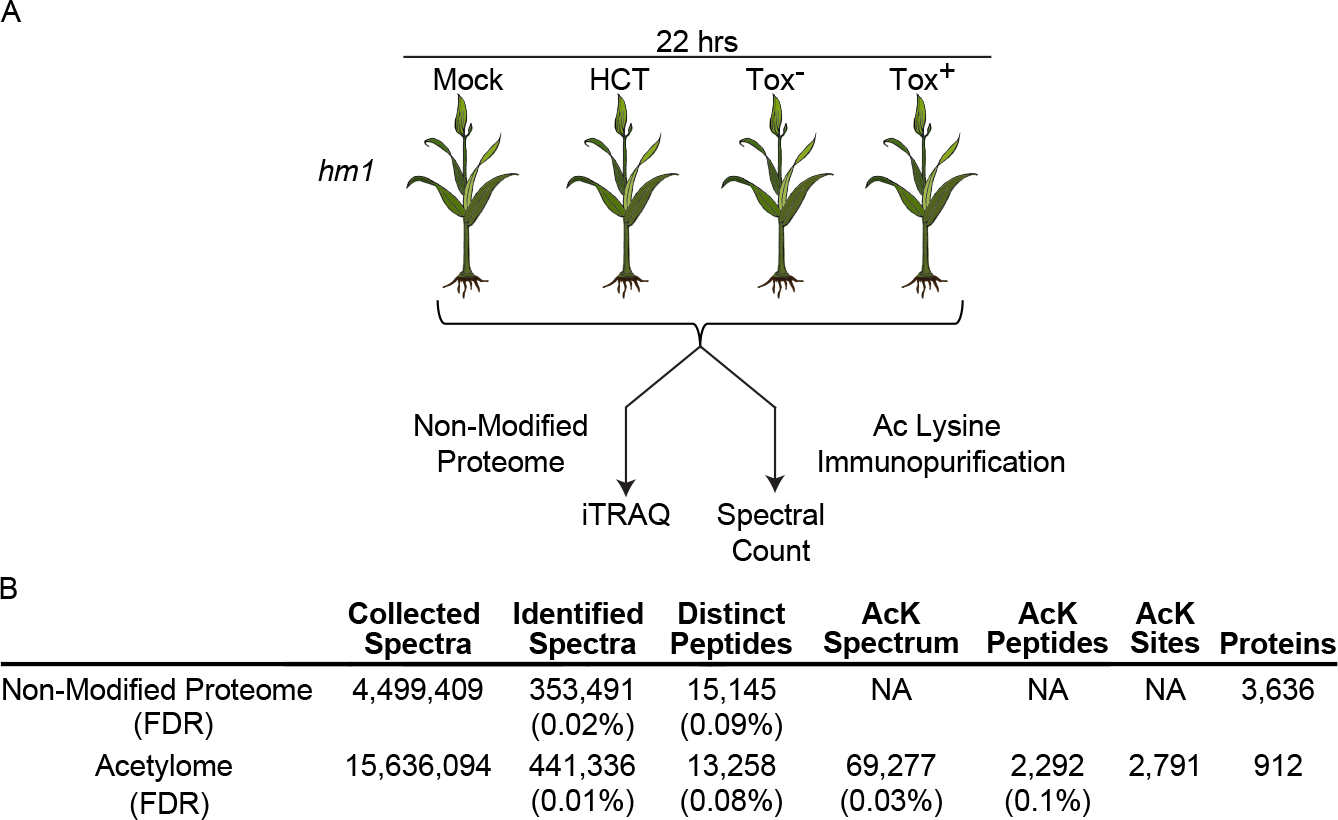
Overview of treatments and proteome profiling. A) Susceptible *Hm1A* plants were exposed to mock, 100 µM HC-toxin, HC-toxin deficient (Tox-), or HC-toxin producing (Tox+) strains of *Cochlibolus carbonum* race 1. For each condition four biological replicates were collected 22 hrs post treatment to quantify protein abundance (iTRAQ) and acetylation levels (spectral counting). B) Summary of sampled spectra, peptides, acetylated peptides, and identified proteins.

### Acetylation is a Global Protein Modification

To gain insight into the composition of the maize acetylome we examined the distribution of acetylated proteins within MapMan functional categories (36). While acetylation is now regarded as a widespread post-translational modification we were surprised that 32 of the 35 major MapMan bins (i.e. functional categories) contained acetylated proteins (Fig. 2 and Dataset S2). Bins that did not contain acetylated proteins are the sparsely populated bins “fermentation”, “polyamine metabolism”, and “microRNA, natural antisense” comprised of 59, 41 and 3 proteins, respectively, which is a small percentage of the 63,542 MapMan annotated proteins. Therefore, it is possible that sampling additional tissues and/or deeper acetylome profiling may reveal that proteins in these functional categories are also acetylated.

We found that the third largest bin is “RNA”, which contains proteins annotated to be involved in RNA splicing and transcriptional regulation (Fig. 2). While earlier studies have described acetylated plant transcription factors the prevalence of this functional category is greater than previously reported (7). Potentially, the increased identification of transcriptional regulatory proteins is due to increased acetylome depth.

We observed acetylation of a wide-range of “classical” maize genes (Dataset S2), which are a set of maize genes that have been intensively studied (37). For example, photosynthetic proteins including light harvesting proteins (LHCB3, LHCB7, LHCB9) as well as the small and large subunits of Rubisco (RBCL, SSU3) are acetylated, which is consistent with the acetylation of homologous proteins in Arabidopsis (7, 8). We observed acetylation of phosphoenolpyruvate carboxylase (PEP1, PEP4) proteins. Additionally, oxylipin signaling proteins lipoxygenase (LOX6; (38) and 12-oxo-phytodienoic acid reductase (OPR1; (39) were detected as acetylated proteins. We also identified acetylation of starch biosynthetic enzymes including sucrose syntase1 (SUS1), UDP-glucose pyrophosphorylase1 (UGP1), ADP Glucose Pyrophosphorylase Small Subunit Leaf1(AGPSLZM), and starch branching enzyme 3 (SBE3) (40–42). Finally, genes involved in transcriptional regulation such as ramosa1 enhancer locus 2 (REL2), a transcriptional co-repressor (43), are acetylated. Taken together these data demonstrate acetylation of a diverse set of proteins and opens the door for new lines of investigation into the regulation of intensively studied maize proteins.

**Figure 2.**
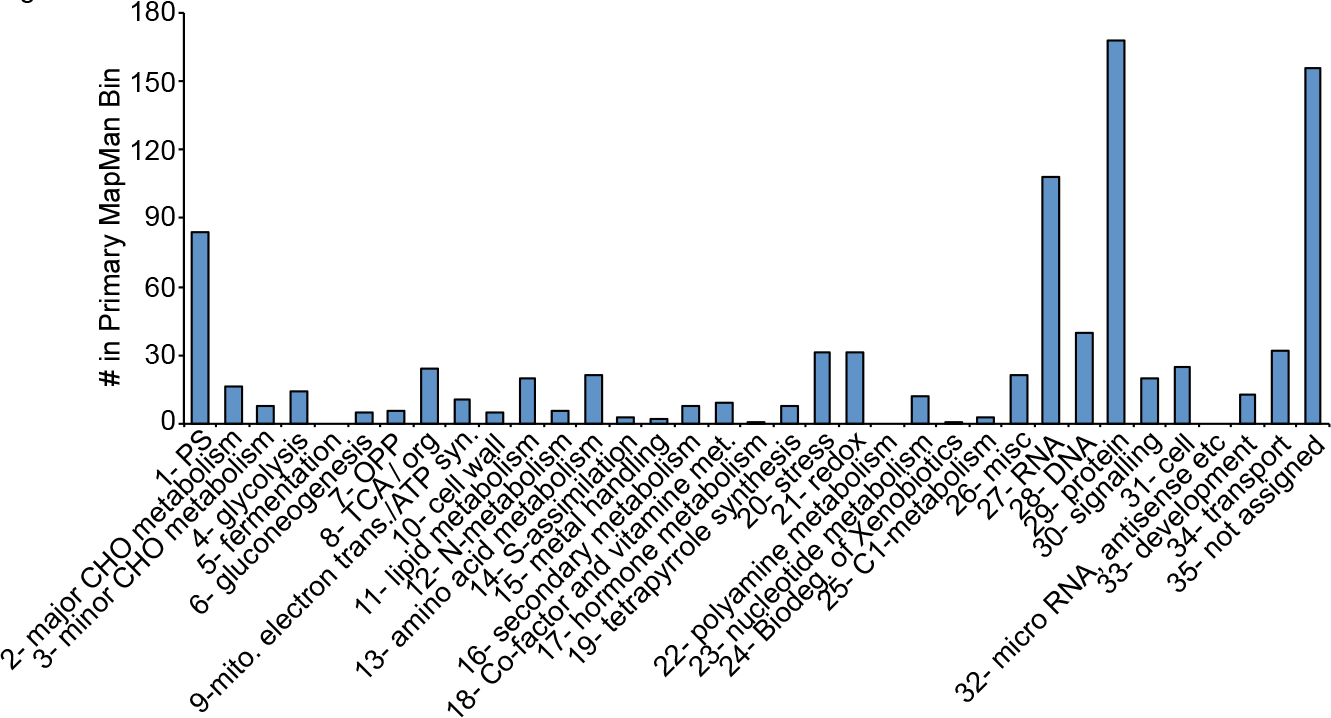
Diverse maize proteins are acetylated. Proteins from 32 out of 35 Major MapMan bins are acetylated.

### Pathogen Infection Alters Histone and Non-Histone Protein Acetylation

To better understand how pathogen infection remodels the host proteome we quantified non-modified protein abundance and acetylation levels. We first verified our acetylation measurements using a commercial antibody that recognizes tetra acetylated histone 4 (H4K5/8/12/16). We found that the H4 tetra acetylation pattern quantified using mass spectrometry matched the pattern determined using western blotting (Fig. 3A&B and Dataset S2). While it has previously been shown that H4 acetylation increases following HC-toxin treatment the specific sites of acetylation were not previously identified (22, 23). Our results demonstrate that H4 tetra acetylation induced by HC-toxin or Tox^+^ infection occurs on H4 lysine residues 5,8,12 and 16.

Globally, we determined that 171 and 116 proteins increased or decreased in abundance following treatment with HC-toxin or infection by *C. carbonum* strains (Fig. 3C and Dataset S1). Additionally, 62 acetylated peptides (155 sites) increased following treatment while only 9 acetylated peptides (12 sites) decreased (Fig. 3D&E). Furthermore, the majority of hyperacetylation events occurred in either the HC-toxin or Tox^+^ treatments but not in response to Tox^-^ infection (Fig. 3D&E). This observation is consistent with HC-toxin functioning as an HDACi to promote protein hyperacetylation.

To gain insight into how HC-toxin induced hyperacetylation promotes *C. carbonum* virulence we performed Gene Ontology (GO) overrepresentation analyses. First, we examined GO categories overrepresented among the acetylated peptides that were altered in abundance following HC-toxin treatment or infection with *C. carbonum* strains (Dataset S3). Strikingly, there are numerous GO categories related to transcriptional regulation over-represented in the HC-toxin and Tox^+^ treatments (Fig. 3F). However, there were no GO categories associated with transcriptional regulation among the acetylated peptides that respond to Tox^-^ infection. Furthermore, defense related GO terms were only present among the Tox^+^ responding acetylated peptides. Taken together these data suggest that virulent *C. carbonum* utilizes HC-toxin to reprogram the transcriptional response to infection resulting in an inappropriate defense response. Furthermore, another non-mutually exclusive possibility is that acetylation of defense proteins by Tox^+^ infection results in inactivation of their function and suppresses host defense.

**Figure 3.**
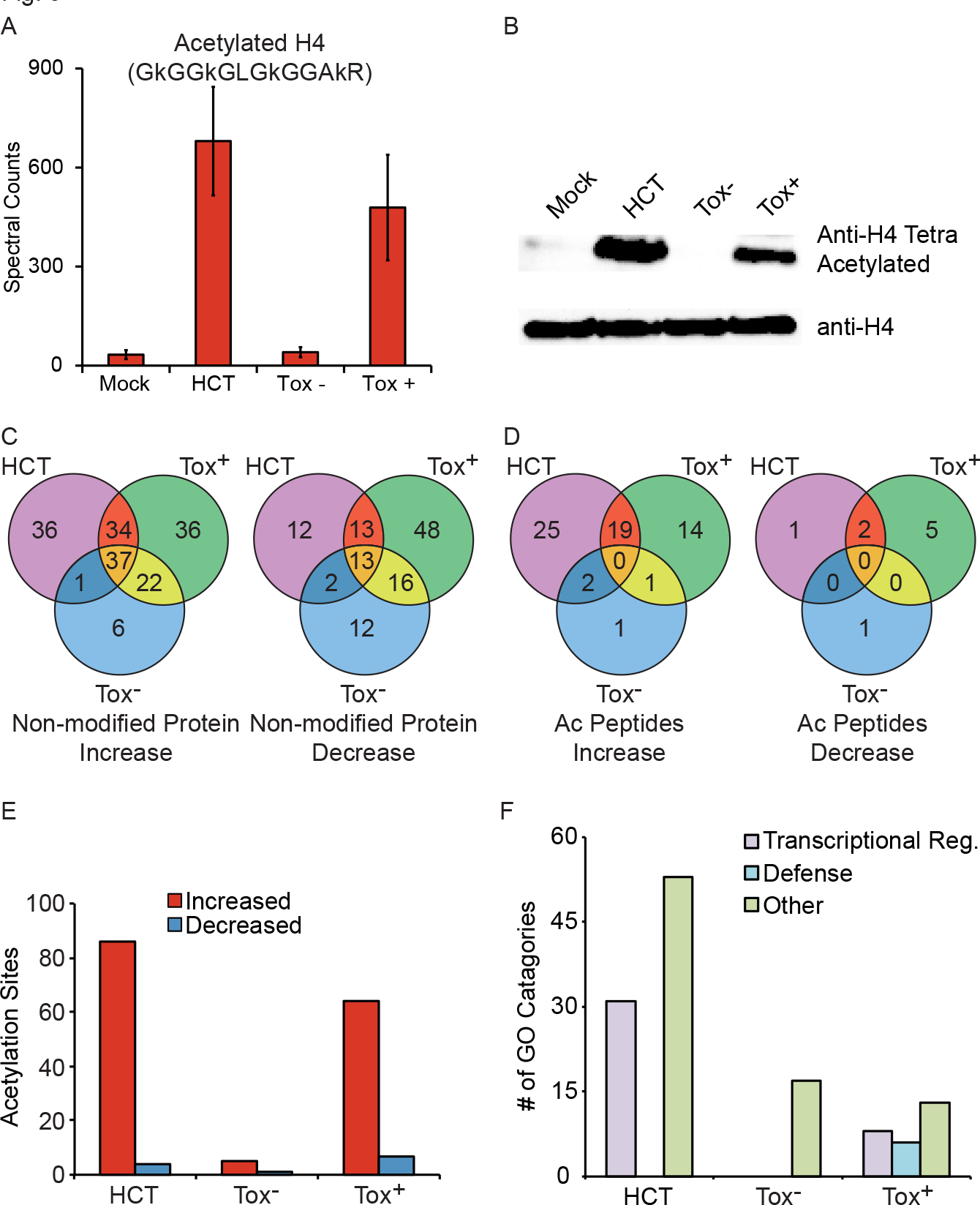
Dynamics of protein abundance and acetylation during immune response. A) Spectral counts of the peptide corresponding to H4 tetra acetylation of lysines 5,8,12 and 16. Data are means of 4 independent biological replicates ± SEM. B) Western blot confirmation of H4 tetra acetylation quantified by proteomics. C) Overlap of protein abundance changes in response to treatments. D) Overlap of acetylated peptides that change in response to treatments. E) Number of acetylation sites that are altered, relative to mock, by HC-toxin, Tox^-^ or Tox^+^ treatment. F) Number of GO categories related to Transcriptional Response, Defense, any other process (“Other”) that are over-represented in HC-toxin, Tox^-^, or Tox^+^ treatment.

### Tryptophan Biosynthetic Proteins are Induced by HC-toxin

To uncover biological processes that may be responding to HC-toxin induced acetylation of transcriptional regulators we used our protein abundance data as a proxy for transcript levels. We identified GO enrichment for a number of biosynthetic processes that are specific to HC-toxin and Tox+ treatments (Fig. S1). We were intrigued by enrichment of GO terms related to tryptophan biosynthesis in HC-toxin and Tox^+^ treated plants and examined these genes further. This analysis revealed that proteins required for every step in the biosynthesis of tryptophan from chorismate were increased in HC-toxin and/or Tox^+^ treatments (Fig. 4). The only protein that peaked following Tox^-^ inoculation was a single isoform of indole-3-glycerol-phosphate synthase. While further research is necessary to verify that HC-toxin transcriptionally activates expression of tryptophan genes, these findings suggest that *C. carbonum* alters tryptophan biosynthesis in order to promote infection.

**Figure 4.**
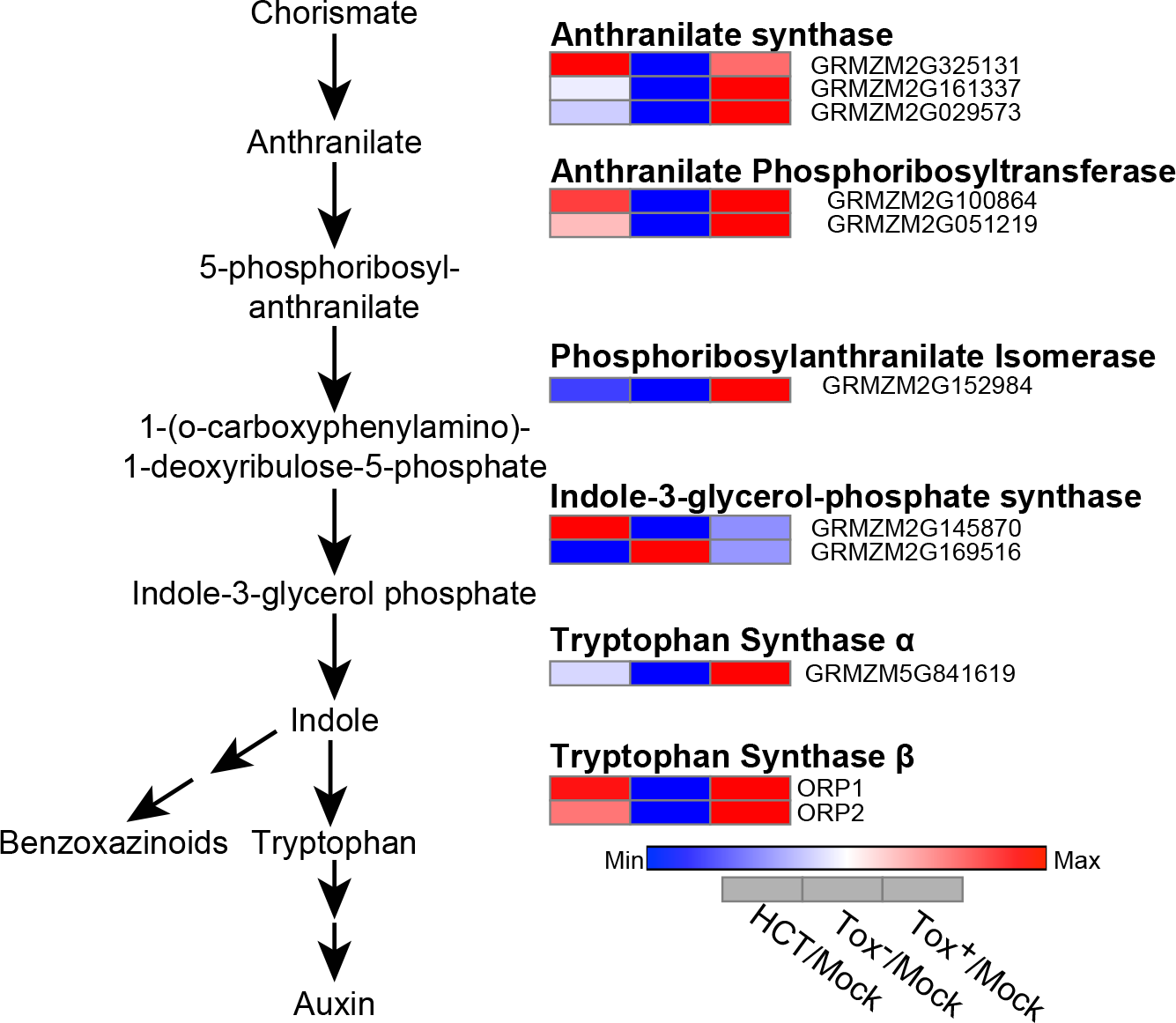
HC-toxin increases the levels of tryptophan biosynthetic proteins. All detected proteins related to each step in tryptophan biosynthesis are shown. Heatmaps represent the relative abundance of each protein following treatment by HC-toxin, Tox^-^, or Tox^+^ relative to mock treatment.

## DISCUSSION

In this study we used global mass spectrometry-based acetylome profiling in maize to identify over 2,700 acetylation sites arising from 912 proteins following exposure to HC toxin, a naturally occurring HDACi. These acetylated proteins represent a wide-range of functional categories suggesting that this post-translational modification can regulate diverse biological processes. Notably, we discovered that many well-characterized maize proteins are acetylated, including proteins responsible for major commercial traits including starch and oil biosynthesis. These novel descriptions of acetylation sites enable new approaches to study the potential regulation of well-characterized proteins and agronomically important traits.

Pathogen infection has been shown to induce non-histone protein acetylation and to alter host immunity (21, 28–32). However, the induced non-histone protein acetylation events previously identified are a result of pathogen effector molecules functioning as acetyltransferase enzymes that act on host proteins. Here, through the application of the exogenous HDACi HC-toxin and *C. carbonum* strains we demonstrate that the activity of plant-encoded histone deacetylase enzymes can be modulated resulting in both histone and non-histone protein hyperacetylation. Thus, endogenous plant enzymes can modulate the acetylation of host non-histone proteins during immune signaling.

We observed that proteins associated with transcriptional regulation are hyperacetylated specifically in response to HC-toxin and Tox^+^ treatment. These transcriptional regulatory proteins are not limited to histones and include transcription factors, chromatin remodeling enzymes and chromatin modifying enzymes. The role of transcription factors in plant defense is well documented and the importance of both chromatin remodeling and modifying enzymes is now recognized (26, 44, 45). Furthermore, acetylation of these types of transcriptional regulatory proteins is known to both reduce or enhance their function (46–49). This suggests that the observed HC-toxin induced hyperacetylation will result in alteration of the transcriptional response during pathogen infection thereby promoting pathogen virulence either through induction of an inappropriate immune response and/or induction of a suppressor(s) of defense.

Finally, by measuring protein abundance levels we determined that treatment with either HC-toxin or Tox^+^ *C. carbonum* results in increased abundance of metabolism related proteins. We are particularly intrigued by the observed increase in proteins in the tryptophan biosynthesis pathway, which may promote susceptibility through several mechanisms. For example, an upregulation of the tryptophan biosynthesis pathway may result in increased auxin biosynthesis (Fig. 4) thereby reducing plant resistance. Consistently, auxin is known to promote either susceptibility or resistance, depending on the pathogen and host (50–54). In addition, an increase in tryptophan biosynthesis may result in increased levels of benzoxazinoids, which are a class of defensive secondary metabolites found in grass species (55–58). In this scenario benzoxazinoids would represent inappropriate defensive metabolites to *C. carbonum* and their induction would thereby reduce resources necessary for maize to mount an effective defense response. As we did not directly measure auxin or benzoxazinoid biosynthetic proteins or the metabolites themselves future work will need to address whether either metabolite directly functions in promoting *C. carbonum* virulence.

## MATERIALS and METHODS

### Plant Material

*Zea mays* plants in which the *Hm1A* (59) allele was introgressed into the B73 inbred were used for all experiments. Plants were grown in a growth chamber in a 16h light/ 8h dark photoperiod at temperatures of 28C (day) and 24C (night). Leaves 2,3 and 4 of fifteen-day-old plants were sprayed mid-day till runoff with either mock (0.1% Tween-20), 100 µM HC-toxin (Sigma) or 400,000 spores/ml of HC-toxin deficient (Tox-) or HC-toxin producing (Tox+) strains of *Cochlibolus carbonum* race 1. Following treatment plants were bagged to increase humidity and placed back in the growth chamber for 22 hrs at which point tissue was collected and flash-frozen. Four biological replicates were used for each treatment.

### Proteomics

Peptide preparation, and non-modified protein profiling by mass spectrometry are based on previously described methods (42, 60). For acetylated peptide enrichment, 2 mg of anti-acetyl-lysine antibody immobilized on agarose beads (ImmuneChem-ICP0388) was added to ~10 mg of maize peptides in 50 mM Tris-HCL pH 7.4. Samples for each biological replicate series (ex. Mock_Rep1, HC-toxin_Rep1, Tox-_Rep1 and Tox+_Rep1) were processed in parallel. The antibody-peptide mixture was incubated for 1 hr with rotation at 4C on a 0.2 uM centrifugal device (Microsep MCPM02C68). Following incubation the beads were washed 3 times with 50 mM Tris-HCL pH 7.4 and then acetyl peptides were eluted using 1.5 ml of 0.1% trifluoracetic acid. The antibody-conjugated beads were washed 2 times with 50 mM Tris-HCL pH 7.4 and then used for a second round of immunoprecipitation of the same sample (i.e. the original flow-through). The 2 enrichments for a given sample were then pooled and passed over a Sep-Pak C18 column (Waters WAT054960) prior to liquid chromatography-mass spectrometry (LC-MS/MS). Full proteomic methods are detailed in SI Materials and Methods.

## FOOTNOTES

Data deposition: The raw spectra for the proteome data have been deposited in the Mass spectrometry Interactive Virtual Environment (MassIVE) repository (massive.ucsd.edu/ProteoSAFe/static/massive.jsp) under the accession ID MSV000079681.

Author contributions: JWW, ZS and SPB designed research; JWW, ZS and MRM

Performed research; JWW and ZS analyzed data; JWW wrote the paper.

## ACKNOWLEDGMENTS

We thank Dior Kelly for critical and thoughtful comments regarding this manuscript, and Gurmukh Johal for seeds of B73 with introgressed Hm1A. This work was supported by a new-faculty start-up grant to JW from Iowa State University, NIH NRSA F32GM096707 to JWW, and National Science Foundation Grant 0924023 to SPB.

